# Time-kill curve analysis and pharmacodynamic functions for *in vitro* evaluation of antimicrobials against *Neisseria gonorrhoeae*

**DOI:** 10.1101/028506

**Authors:** Sunniva Foerster, Magnus Unemo, Lucy J. Hathaway, Nicola Low, Christian L. Althaus

## Abstract

Gonorrhea is a sexually transmitted infection caused by the Gram-negative bacterium *Neisseria gonorrhoeae*. Resistance to first-line empirical monotherapy has emerged, so robust methods are needed to appropriately evaluate the activity of existing and novel antimicrobials against the bacterium. Pharmacodynamic functions, which describe the relationship between the concentration of antimicrobials and the bacterial net growth rate, provide more detailed information than the MIC only. In this study, a novel standardized *in vitro* time-kill curve assay was developed. The assay was validated using five World Health Organization *N. gonorrhoeae* reference strains and various concentrations of ciprofloxacin, and then the activity of nine antimicrobials with different target mechanisms were examined against a highly susceptible clinical wild type isolate (cultured in 1964). From the time-kill curves, the bacterial net growth rates at each antimicrobial concentration were estimated. Finally, a pharmacodynamic function was fitted to the data, resulting in four parameters that describe the pharmacodynamic properties of each antimicrobial. Ciprofloxacin resistance determinants shifted the pharmacodynamic MIC (zMIC) and attenuated the bactericidal effect at antimicrobial concentrations above the zMIC. Ciprofloxacin, spectinomycin and gentamicin had the strongest bactericidal effect during the first six hours of the assay. Only tetracycline and chloramphenicol showed a purely bacteriostatic effect. The pharmacodynamic functions differed between antimicrobials, showing that the effect of the drugs at concentrations below and above the MIC vary widely. In conclusion, *N. gonorrhoeae* time-kill curve experiments analyzed with pharmacodynamic functions have potential for *in vitro* evaluation of new and existing antimicrobials and dosing strategies to treat gonorrhea.

---

Antimicrobial resistance in *Neisseria gonorrhoeae* is a major public health problem. Strains of *N. gonorrhoeae* have developed resistance to all antimicrobials introduced for treatment and rare strains have been classified as superbugs. Clinical resistance to the last option for empirical antimicrobial monotherapy, ceftriaxone, was firstly described in 2009 (1, 2). Currently, treatment recommendations for gonorrhea and prediction of the efficacy of antimicrobials mainly rely on a single measurement: the MIC of the antimicrobial. However, antimicrobials that have different modes of action and lead to different treatment outcomes can have identical MICs (2). A better understanding of the *in vitro* pharmacodynamic properties of antimicrobials could be used to optimize dosing strategies and help prevent treatment failures (3).

Regoes et al. (4) introduced the concept of pharmacodynamic functions to study the relationship between bacterial net growth rates and the concentrations of antimicrobials, based on analyses of time-kill curves for a single laboratory strain of *Escherichia coli.* Mathematically, the pharmacodynamic function is based on a Hill function and characterized by four parameters: the maximal bacterial growth rate in the absence of antimicrobial (*ψ*_max_), the minimal bacterial growth rate at high concentrations of antimicrobial (*ψ*_min_), the Hill coefficient (*κ*), and the pharmacodynamic MIC (zMIC) (Figure 1). Information about the effects of antimicrobials at concentrations below and above the MIC is particularly valuable for pathogens like *N. gonorrhoeae*, for which there are limited data about the pharmacokinetic and pharmacodynamic effects of many antimicrobials. Furthermore, the pharmacodynamic properties of novel antimicrobials with known and unknown targets could be evaluated and directly compared to a set of mechanistically well-understood compounds.

**FIG 1.**
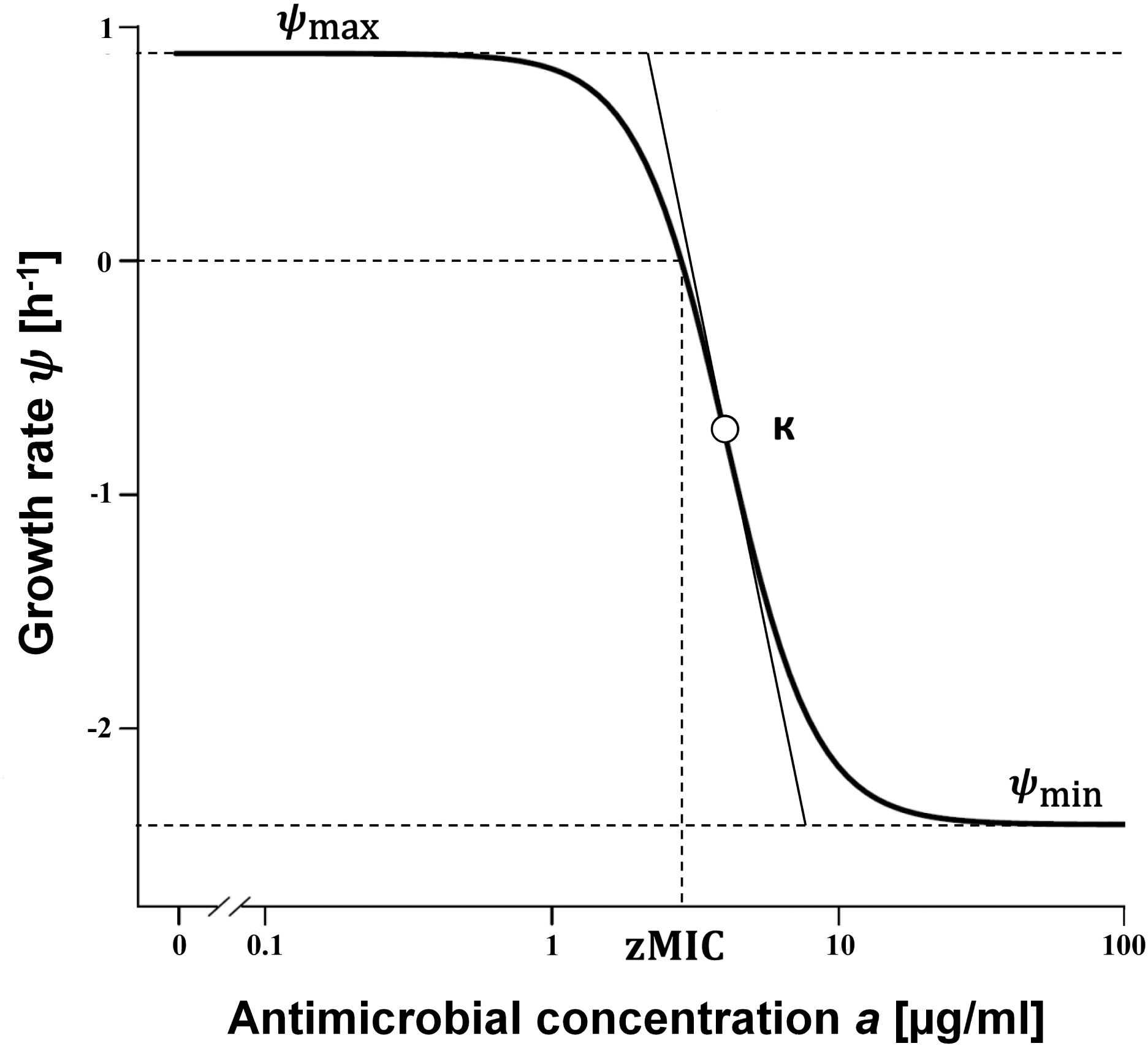
The pharmacodynamics function and four parameters is shown as described in (4). The bacterial growth rates (*ψ*) in response to each antimicrobial concentration are estimated from time-kill data with linear regression. The maximal bacterial growth rate *ψ*_max_, the minimal bacterial net growth rate at high concentrations of antimicrobial *ψ*_min_, the pharmacodynamic MIC (zMIC) and the Hill coefficient *κ* are shown and define the shape of the curve.

There is no standardized and quality assured procedure for time-kill curve analysis of the fastidious obligate human pathogen *N. gonorrhoeae.* Published time-kill protocols for *N. gonorrhoeae* (5–7) are not generalizable, owing to the highly divergent growth requirements of different strains and interpretation of results generally relies on qualitative expert judgement. To study a wide range of *N. gonorrhoeae* strains, growth in the absence of antimicrobials must be consistent and comparable and bacterial growth phases at the time of exposure to antimicrobial need to be synchronized in early to mid-log phase.

In this study, a standardized *in vitro* time-kill curve assay for *N. gonorrhoeae* was developed using Graver-Wade (GW) medium. GW medium is a chemically defined, nutritious, liquid medium that supports growth of a wide range of *N. gonorrhoeae* auxotypes and clinical isolates starting from very low inocula (8). The novel time-kill curve assay was validated on five World Health Organization *N. gonorrhoeae* reference strains with fluoroquinolone resistance determinants. A highly susceptible *N. gonorrhoeae* isolate (DOGK18, 1964, Denmark) was subsequently studied in detail and time-kill curve analysis performed for nine antimicrobials that have been or currently are used to treat gonorrhea. In a second step, we obtained the pharmacodynamic functions for each antimicrobial from the *in vitro* time-kill data and studied their pharmacodynamic properties against *N. gonorrhoeae.*

## MATERIALS AND METHODS

*Neisseria gonorrhoeae* **isolates and media**. The five international *N. gonorrhoeae* reference strains WHO G, WHO K, WHO L, WHO M, and WHO N with different ciprofloxacin conferring mutations in *gyrA, parC* and *parE* (9) and a clinical antimicrobial susceptible ‘wild-type’ isolate cultured in 1964 in Denmark (DOGK18), were studied. Isolates were cultured, from frozen stocks (-70°C), on GCAGP agar plates (3.6% Difco GC Medium Base agar [BD, Diagnostics, Sparks, MD, USA] supplemented with 1% haemoglobin [BD, Diagnostics], 1% IsoVitalex [BD, Diagnostics] and 10% horse serum) for 18-20 hours at 37°C in a humid 5% CO2-enriched atmosphere. Gonococcal colonies were subcultured once more on GCAGP agar for 18-20 hours at 37°C in a humid 5% CO_2_-enriched atmosphere, before being transferred to the liquid sterile GW medium, prepared as earlier described (4), for growth curve and time-kill experiments.

**Viable cell counts.** Bacterial viability was measured using a modified Miles and Misra method as previously described (10). Growing bacteria were removed from 96-well plates at specified time points using a multichannel pipette and diluted in sterile phosphate buffered saline (PBS) in six subsequent 1:10 dilutions (20 μl culture in 180 μl diluent). Ten μl droplets of each dilution were spotted on GCRAP (3.6% Difco GC Medium Base agar [BD, Diagnostics] supplemented with 1% hemoglobin [BD, Diagnostics] and 1% IsoVitalex [BD, Diagnostics]). GCRAP plates were dried with open lid in a sterile environment for 30-60 minutes prior to their usage. After drying the droplets (approximately 5-10 minutes), plates were incubated for 24 hours at 37°C in a humid 5% CO_2_-enriched atmosphere. For every concentration and time point, colonies were counted for the first dilution that resulted in a countable range of 3-30 colonies and the CFU/ml calculated.

**Growth curves.** Prior to growth curve experiments strains were subcultured on Chocolate agar PolyViteX (Biomerieux). A 0.5 McFarland inoculum was prepared and diluted to 100 CFU/ml (1:10^6^) in GW Medium (35°C). A volume of 100 μl diluted bacteria per well was transferred to Sarstedt round bottom 96 well plates. The plates were tightly sealed and bacteria were grown shaking at 100 rpm at 35°C in a humid 5% CO_2_-enriched atmosphere. Bacterial growth was monitored over a time-course of sixty hours (0, 2, 4, 6, 8, 10, 12, 20, 22, 24, 26, 28, 30, 32, 34, 40, 44, 48, 60 hours). For every sampled time point, the content of one well was removed and viable counts determined (10). Growth curves were analyzed by plotting the log CFU/ml against the time and fitting a Gompertz growth model to the data as implemented in the package *cellGrowth* (11, 12). Only lag, log and stationary phases were included in the analysis and the decline phase excluded.

**Time-kill assay.** Time-kill curve analyses were performed by culturing *N. gonorrhoeae* in GW medium (4), in the presence of eleven antimicrobial-concentrations in doubling dilutions ranging from 0.016×MIC to 16×MIC. For DOGK18, the MICs were determined prior to the experiment using Etest (bioMérieux, Marcy l’Etoile, France) according to the manufacturer’s instructions. For all other strains, previously published MIC values were used (9). The antimicrobials examined were ciprofloxacin (Sigma Aldrich, China), gentamicin (Sigma Aldrich, Israel), spectinomycin (Sigma Aldrich, Israel), azithromycin (Sigma Aldrich, USA), benzylpenicillin (Sigma Aldrich, USA), ceftriaxone (Sigma Aldrich, Israel), cefixime (European pharmacopeia reference standard, France), chloramphenicol (Sigma Aldrich, China) and tetracycline (Sigma Aldrich, China). Growth curves were initially performed to confirm that all strains would reach a stable early- to mid-log phase after four hours of pre-incubation in antimicrobial-free GW medium. Subsequently, a 0.5 McFarland inoculum of *N. gonorrhoeae* was prepared in sterile PBS from cultures grown on GCAGP agar plates for 18–20 hours at 37°C in a humid 5% CO2-enriched atmosphere. For each strain, 30 μl of the inoculum was diluted in 15 ml pre-warmed (37°C) antimicrobial-free GW medium and 90 μl per well was dispersed in round bottom 96-well Sarstedt microtiter plates. The plates were pre-incubated for 4 h shaking at 150 rpm, 35°C in a humid 5% CO_2_-enriched atmosphere. To each well containing 90 μl of pre-incubated bacteria, 10 μl of one of the antimicrobial concentrations (or PBS) was added, resulting in eight identical rows (one row for each time-point) containing bacteria exposed to eleven different antimicrobial concentrations and one untreated control.

**Estimating bacterial growth rates.** The bacterial net growth rates (*ψ*) were determined from changes in the density of viable bacteria (CFU/ml) during the first six hours of the time-kill experiments. The bacterial populations were assumed to grow or die at a constant rate, resulting in an exponential increase or decrease in bacterial density:

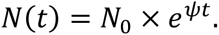

The net growth rate was estimated as the coefficient of a linear regression from the logarithm of the colony counts. Maximum likelihood estimation was used to account for the censored data (values below the limit of detection of 100 CFU/ml). The geometric mean of all measurements at zero hours for a given antimicrobial as the first data point, was used. From the growth rate, the bacterial doubling time can be calculated as follows:

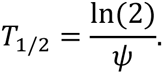

**Pharmacodynamic function.** A pharmacodynamic function (Fig. 1) describes the relationship between bacterial net growth rates (*ψ*) and the concentration of an antimicrobial (*a*) (4):

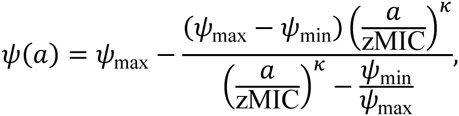

where *ψ*_max_ is to the maximal bacterial growth rate in the absence of antimicrobial and *ψ*_min_ is the minimal bacterial net growth rate at high concentrations of antimicrobial. zMIC is the pharmacodynamic MIC where the bacterial growth rate is zero (*ψ*(zMIC) = 0). *κ* denotes the Hill coefficient, which describes the steepness of the sigmoid relationship between bacterial growth and antimicrobial concentration. For each antimicrobial, four parameters of the pharmacodynamic function were estimated using a self-starter function, implemented in the package *drc* (13) for the R software environment for statistical computing (14). All figures can be reproduced with R code and data from the following GitHub repository: https://github.com/sunnivas/PDfunction.

## RESULTS

**Growth of *N. gonorrhoeae***. Growth curves for the five different WHO *N. gonorrhoeae* reference strains (Fig. S1 supplemental material) confirmed that growth was well supported in GW medium. All strains could be grown from a starting inoculum of less than 10^3^ CFU/ml and had a lag phase of less than five hours. The stationary phase lasted until 36 hours for all strains, followed by a steep declination phase. Growth was similar for all strains with WHO L the only strain that had a slightly slower growth.

**Time-kill curves.** Time-kill curves for ciprofloxacin using the WHO reference strains WHO G (MIC = 0.125 μg/ml), WHO K (MIC > 32 μg/ml), WHO L (MIC > 32 μg/ml), WHO M (MIC = 2 μg/ml), WHO N (MIC = 4 μg/ml) and DOGK18 (MIC = 0.008 μg/ml) are shown in Fig. 2. Ciprofloxacin induced a bactericidal effect in all six strains, but the onset of the bactericidal activity depended on the concentration of the antimicrobial and differed between strains. All strains with the exception of WHO M and WHO N were killed to below the limit of detection (100 CFU/mL) at the highest antimicrobial concentration (16×MIC). The susceptible DOGK18 strain experienced the most rapid killing during the first hour at high antimicrobial concentrations. For WHO G and WHO M, the bactericidal activity decreased during the six hours of the assay.

**FIG 2.**
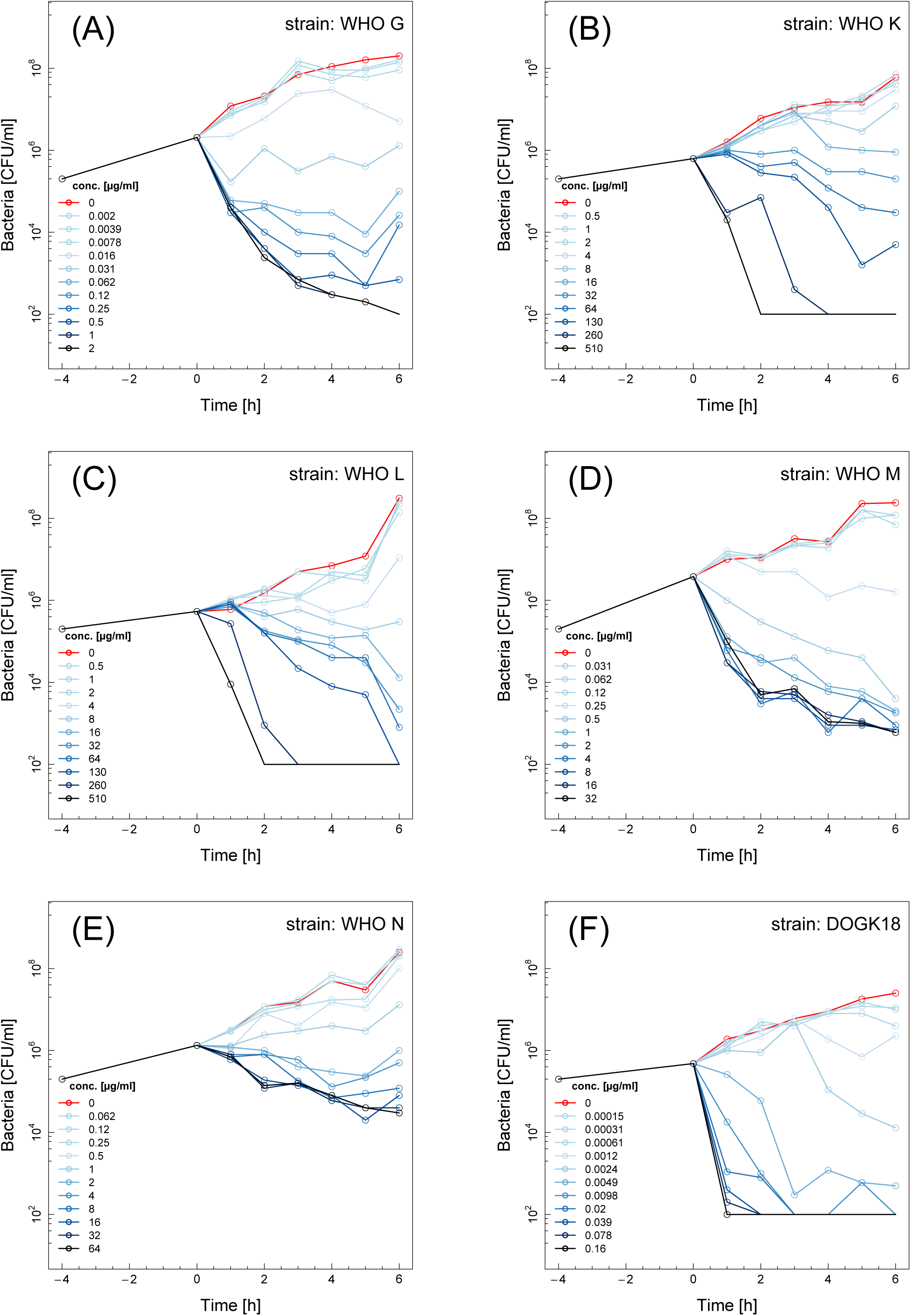
Time-kill curves for ciprofloxacin and six different *Neisseria gonorrhoeae* strains: WHO G (A), WHO K (B), WHO L (C), WHO M (D), WHO N (E) and DOGK18 (F). Controls without antimicrobials are shown in red. Twelve doubling dilutions are plotted, the highest concentration (black line) corresponds to 16× MIC as measured with Etest. The limit of detection in the assay was 100 CFU/ml. Antimicrobial was added at time 0 h.

Time-kill curves for eight antimicrobials were made (spectinomycin, gentamicin, azithromycin, benzylpenicillin, ciprofloxacin, ceftriaxone, cefixime, chloramphenicol and tetracycline) using the highly antimicrobial susceptible DOGK18 strain (Fig. 3). Similar to the effect of ciprofloxacin (Fig. 2F), gentamicin and spectinomycin exhibited rapid killing during the first two hours of the assay for concentrations above MIC. Cefixime and ceftriaxone showed little effect from 0 hours to 3 hours, however, after that growth rate decreased rapidly. For benzylpenicillin and azithromycin, at concentrations above MIC, the killing started after one hour and decreased rapidly at later time points. The time-kill curves for tetracycline and chloramphenicol looked similar with almost no killing of bacteria within the assay time of four hours. Chloramphenicol showed a weak bactericidal effect at the highest antimicrobial concentration (Fig. 3).

**FIG 3.**
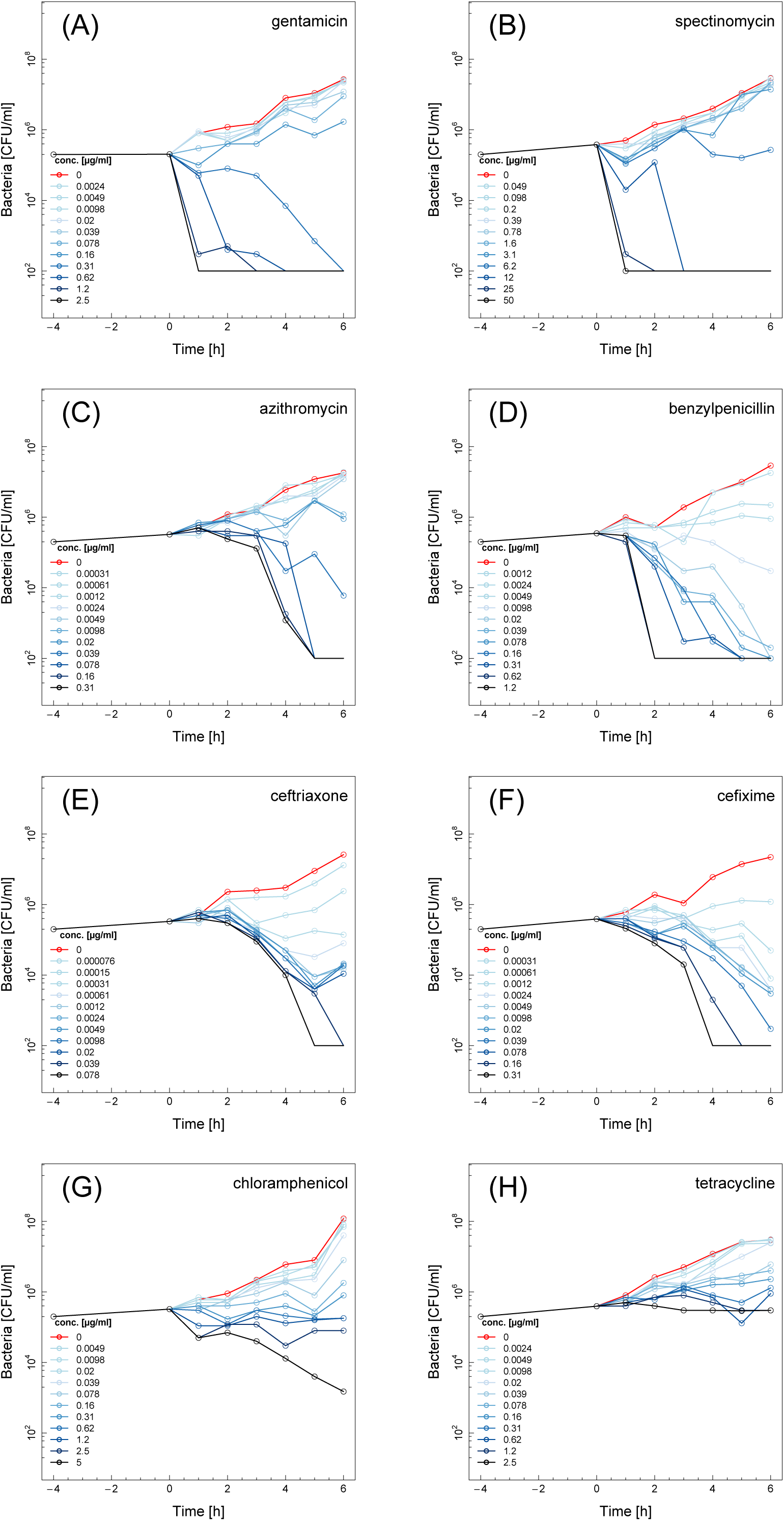
Time-kill curves for the *Neisseria gonorrhoeae* DOGK18 strain using eight different antimicrobials: gentamicin (A), spectinomycin (B), azithromycin (C), benzylpenicillin (D), ceftriaxone (E), cefixime (F), chloramphenicol (G) and tetracycline (H). Twelve doubling dilutions are plotted, the highest concentration (black line) corresponds to 16× MIC as measured with Etest. The limit of detection in the assay was 100 CFU/ml. Antimicrobial was added at time 0 h.

**Pharmacodynamic functions.** The bacterial net growth rates were estimated from the time-kill curves by fitting a linear regression to the logarithm of the colony counts (Fig. 4A). The pharmacodynamic function was then fitted to the estimated growth rates at different antimicrobial concentrations (Fig. 4B, solid line). Generally, the estimated net growth rates at high antimicrobial concentrations reached a lower asymptote (*ψ*_min_). In some cases, an additional drop was observed in the estimated growth rates at very high antimicrobial concentrations (Fig. 4B, dashed line). This phenomenon occurred at antimicrobial concentrations that are likely to be toxic, so those data points were removed before estimating the parameters of the pharmacodynamic function.

**FIG 4.**
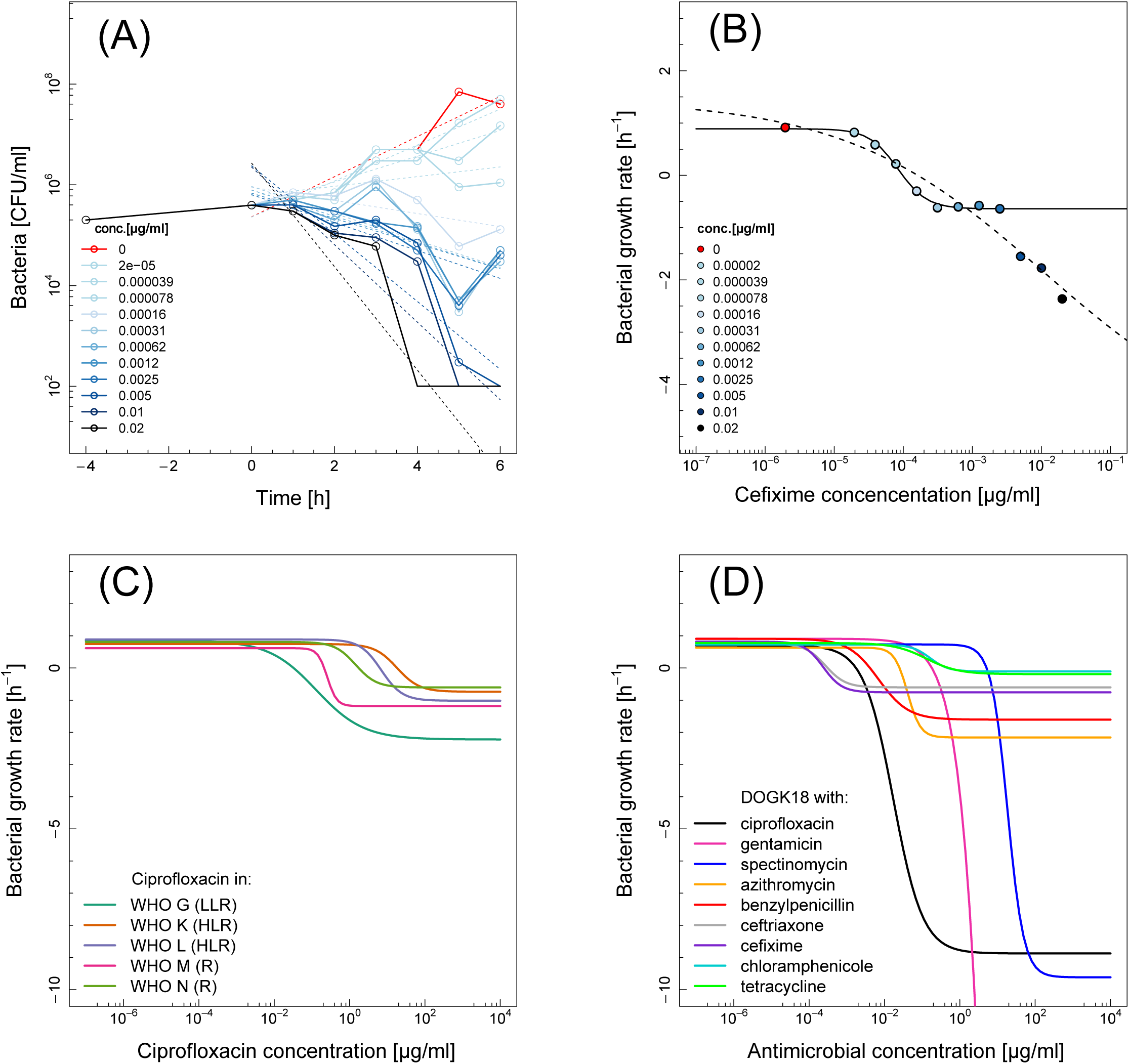
Pharmacodynamic functions for different antimicrobials and *Neisseria gonorrhoeae* strains. (A) Estimating growth rates (cefixime in DOGK18). Dashed lines represent linear regressions of the logarithm of the colony counts at different antimicrobial concentrations. The coefficient of the linear regression corresponds to the net bacterial growth rate (cefixime in DOGK18). (B) Fitting the pharmacodynamic function to estimated growth rates. Points correspond to the estimated net bacterial growth rates at different antimicrobial concentrations. The solid line shows the model fit after removing outliers at high antimicrobial concentrations. The dashed line indicates the model fit including all data points. The growth rate in absence of antimicrobial is shown in red at a concentration that is 10-fold lower than the lowest concentration. (C) Pharmacodynamics functions for ciprofloxacin in six *N. gonorrhoeae* strains (Low level resistance (LLR)=WHO G; High level resistance (HLR)=WHO K, WHO L; Resistance (R)=WHO M, WHO N; and Susceptible (S)= DOGK18) (D) Pharmacodynamic functions for nine different antimicrobials in DOGK18 strain. Each curve is based on the arithmetic mean of the estimated parameters from two independent time-kill experiments (as in Table 1).

The pharmacodynamic functions for the six strains provided information on ciprofloxacin resistance mechanisms (Fig. 4C and Table S3 in the supplemental material). The DOGK18 strain had a low pharmacodynamic MIC (zMIC) and a low net bacterial growth rate at high antimicrobial concentrations (*ψ*_min_), indicating the strong bactericidal effect of ciprofloxacin. The five WHO reference strains showed that the ciprofloxacin resistance determinants not only shifted the zMIC but also resulted in less killing at antimicrobial concentrations above the zMIC (higher *ψ*_min_ compared to DOGK18 strain).

The pharmacodynamic functions for the nine antimicrobials in the DOGK18 strain illustrated the different effects that antimicrobials induce on gonococcal growth of *N. gonorrhoeae* (Fig. 4D and Table 1). The average of the maximal growth rate in the absence of antimicrobials over all experiments was *ψ*_max_ = 0.77 h^−1^ (95% confidence interval [CI]: 0.71-0.84 h^−1^). This corresponds to a bacterial doubling time of *T*_1/2_ = 54 min (95% CI: 49-59 min). Ciprofloxacin, spectinomycin and gentamicin induced the strongest bactericidal effect with *ψ*_min_ < -5 h^−1^. Chloramphenicol and tetracycline exhibited almost no killing within the six hours of the assay (*ψ*_min_ > -0.2 h^−1^). The Hill coefficient *κ* ranged between 1.0 and 2.5. All four parameters of the pharmacodynamic function were very similar for ceftriaxone, cefixime and the bacteriostatic compounds chloramphenicol and tetracycline. Generally, the estimated zMIC was in good agreement with the MIC measured by Etest but there were substantial deviations for benzylpenicillin and cefixime.

**TABLE 1.** Parameter estimates of the pharmacodynamic function for nine antimicrobials in the antimicrobial susceptible *Neisseria gonorrhoeae* strain DOGK18.

| Antimicrobial | Antimicrobial class | $\kappa^a$ | $\psi_{\min} (\text{h}^{-1})^a$ | $\psi_{\max} (\text{h}^{-1})^a$ | zMIC ($\mu\text{g/ml}$)<sup>a</sup> | MIC ($\mu\text{g/ml}$)<sup>b</sup> |
| --- | --- | --- | --- | --- | --- | --- |
| Ciprofloxacin | Fluoroquinolone | 1.1±0.1 | -8.9±2.2 | 0.7±0.4 | 0.002±0.0001 | <0.004 |
| Gentamicin | Aminoglycoside | 2.0±0.3 | -107.4±140.6 | 0.9±0.07 | 0.2±0.04 | 1 |
| Spectinomycin | Aminocyclitol | 1.0±0.6 | -9.6±1 | 0.7±0.03 | 5±0.7 | 4 |
| Azithromycin | Macrolide | 2.5±0.2 | -2.2±0.1 | 0.6±0.05 | 0.3±0.03 | 0.19 |
| Benzylpenicillin | β-lactam | 1.1±0.1 | -1.6±0.6 | 0.9±0.2 | 0.004±0.002 | 0.032 |
| Ceftriaxone | β-lactam | 1.6±0.1 | -0.6±0.2 | 0.8±0.07 | 0.0003±0.0001 | <0.002 |
| Cefixime | β-lactam | 1.7±0.5 | -0.8±0.2 | 0.8±0.1 | 0.0002±0.0002 | 0.016 |
| Chloramphenicol | Chloramphenicol | 1.8±0.4 | -0.1±0.01 | 0.7±0.2 | 0.5±0.1 | 0.19 |
| Tetracycline | Tetracycline | 1.0±0.2 | -0.2±0.08 | 0.8±0.07 | 0.5±0.3 | 0.125 |

<sup>a</sup>Estimates are given as arithmetic means and standard deviations from two independent experiments. Parameter estimates for each individual experiment are given in Table S2 of the supplemental material.
<sup>b</sup>MIC values measured with Etest in accordance with the manufacturer's instructions.

## DISCUSSION

A robust and reliable method to appropriately evaluate antimicrobial treatment options *in vitro* is one of the tools that is urgently needed to support tackling the problem of antimicrobial resistant *N. gonorrhoeae.* In this study, a standardized *in vitro* time-kill curve assay was developed and the resulting data were used to study pharmacodynamics parameters describing the relationship between the concentration of antimicrobials and the bacterial net growth rate. Our study applied the concept of pharmacodynamic functions (4) to *N. gonorrhoeae* for the first time. To our knowledge, this is also the first study to obtain and compare the pharmacodynamic functions of an antimicrobial in susceptible and resistant strains of the same pathogen species, opening up avenues into phenotypically understanding the effects of different resistance determinants.

The developed time-kill assay worked well for different *N. gonorrhoeae* strains, including highly resistant isolates. Growing the bacteria in microwell plates provided a high throughput and made it possible to study a wider range of antimicrobial concentrations in the same experiment. Growth curves confirmed reproducible growth in absence of antimicrobials, WHO L had a prolonged lag phase and grew slightly slower than the other strains. Therefore in the time-kill assays bacteria were grown for four hours before adding the antimicrobial. The assay time was limited to six hours and growth in absence of antimicrobials highly consistent and exponential for all strains. The five WHO reference strains tested have different combinations of ciprofloxacin resistance-conferring mutations in *gyrA, parC* and *parE.* These mutations resulted in a shift of the pharmacodynamic MIC (zMIC) and reduced the antimicrobial killing at the same time, showing that even high doses of ciprofloxacin had a limited effect on the growth of these strains. The analysis of time-kill data with a pharmacodynamic function confirmed the strong bactericidal effects of ciprofloxacin, gentamicin and spectinomycin. Ciprofloxacin is a prime example of a bactericidal antimicrobial, representing the class of topoisomerase II inhibiting fluoroquinolones (15). Spectinomycin inhibits protein translation through blocking of tRNA translocation, whereas gentamicin induces translational misreading (16–18). While spectinomycin is a well-recognised treatment option for gonorrhea and its bactericidal action has been described previously (19, 20), gentamicin is only recommended first-line treatment for gonorrhea in Malawi, where it is used together with doxycycline in the syndromic management algorithm for urethritis (21). However, gentamicin has been suggested for wider use in the treatment of gonorrhea (22–24). The cell wall inhibiting β-lactam antimicrobials are known to have a time-dependent mode of action (25, 26). Therefore it was not surprising that benzylpenicillin, ceftriaxone and cefixime were characterized by slower killing (−1.6 h^−1^ < *ψ*_min_ < 0.6 h^−1^). Although currently not used for treatment of *N. gonorrhoeae,* chloramphenicol and tetracycline often act as model compounds for bacteriostatic effects (27, 28). These effects were confirmed by the estimates of the net bacterial growth rate at high antimicrobial concentrations that were close to zero.

Regoes et al. (4) hypothesized that the Hill coefficient of the pharmacodynamic function (*κ*) reflects the time and concentration dependency of antimicrobials. For example, ciprofloxacin is thought to act in a concentration-dependent manner and a high value of *κ* would require the antimicrobial concentrations to be at or above the zMIC. Tetracycline was considered a time-dependent antimicrobial which could be characterized by lower values of *κ*. Our estimate for tetracycline was not significantly different from that of the bactericidal ciprofloxacin and was not as low as reported for *E. coli* (4). These discrepancies could reflect differences in time and concentration dependency of antimicrobials in *N. gonorrhoeae* and *E. coli*, different concentration ranges studied (up to 100x MIC) or could be simply due to the properties of the specific strain Regoes et al. (4) studied.

There are some limitations of the methods used in the present study. First, the rapid bactericidal effects of some antimicrobials occurred minutes after the compound was added resulting in bacterial counts below limit of detection at the first time point. These effects can make it challenging to estimate the minimal bacterial net growth rate at high concentrations of antimicrobials (*ψ*_min_), as observed for gentamicin for example. Second, exceedingly high antimicrobial concentrations can kill *N. gonorrhoeae* almost instantaneously. This might explain the observed outliers that deviated from the pharmacodynamic function (Fig. 4B). Third, the time-kill curves appeared to level off over time for bactericidal compounds in susceptible strains. Interestingly, this phenomenon might represent a physiological adaptation to those antimicrobials, often described as persister cell formation (29–34). This non-exponential killing makes it difficult to estimate the net growth rate with linear regression. The clinical relevance of persister cells has been demonstrated for chronic infections such as tuberculosis, cystic fibrosis and infections caused by *Staphylococcus aureus* (35–36). For *N. gonorrhoeae*, however, this phenomenon has not been previously described. Fourth, the proposed method allows appropriate evaluation of antimicrobials against *N. gonorrhoeae in vitro* only.

The *in vitro* pharmacodynamic parameters can provide relative comparisons across different strains and antimicrobials which can be extremely valuable in preclinical studies. However, deriving rational resistance breakpoints for different dosing schedules will have to come from clinical pharmacokinetic and pharmacodynamic (PK/PD) studies that include parameters such as serum concentrations and half-life time of the antimicrobial. For benzylpenicillin, ceftriaxone and cefixime the time of free antimicrobial above the MIC value should be maximized (37–40), suggesting that multiple dose treatment would be a rational strategy. However the longer serum half-life of ceftriaxone (41) compared to other β-lactam antimicrobials suggests that increasing the dose is still efficacious for most *N. gonorrhoeae* strains. Fluoroquinolones and aminoglycosides, which act in a concentration dependent and bactericidal manner should be given as a single high dose (42). This is typically achieved maximizing the AUC/MIC and peak serum concentration/MIC ratio (43-45). Our results suggest that this could be the case for ciprofloxacin, gentamicin and spectinomycin, which were found to be strongly bactericidal. Azithromycin has been described to be bacteriostatic in *Staphylococcus aureus, Streptococcus pneumoniae* or *Haemophilus influenza* (46) but appears to act bactericidal on *Pseudomonas aeruginosa* (47). Our in-vitro pharmacodynamics parameters suggest that a clear classification of azithromycin as bacteriostatic or bactericidal is not possible for *N. gonorrhoeae*.

In summary, the present study shows that evaluation of the parameters of a pharmacodynamic function based on time-kill data can add valuable information beyond that of MIC values for different antimicrobials. The quantitative assessment of pharmacodynamic properties provides a more detailed picture about antimicrobial-induced effects on *N. gonorrhoeae.* The developed time-kill curve assay, in combination with the concept of the pharmacodynamic function, could be used for *in vitro* evaluation of new and existing antimicrobials and the effects of combining antimicrobials against *N. gonorrhoeae.*

## ACKNOWLEDGEMENTS

The present study was funded through an Interdisciplinary PhD (IPhD) project from SystemsX.ch (The Swiss Initiative for Systems Biology), RADAR-Go (RApid Diagnosis of Antibiotic Resistance in Gonorrhoea; funded by the Swiss Platform for Translational Medicine), and the Örebro County Council Research Committee and the Foundation for Medical Research at Örebro University Hospital, Sweden.

---

**Figure S1.**
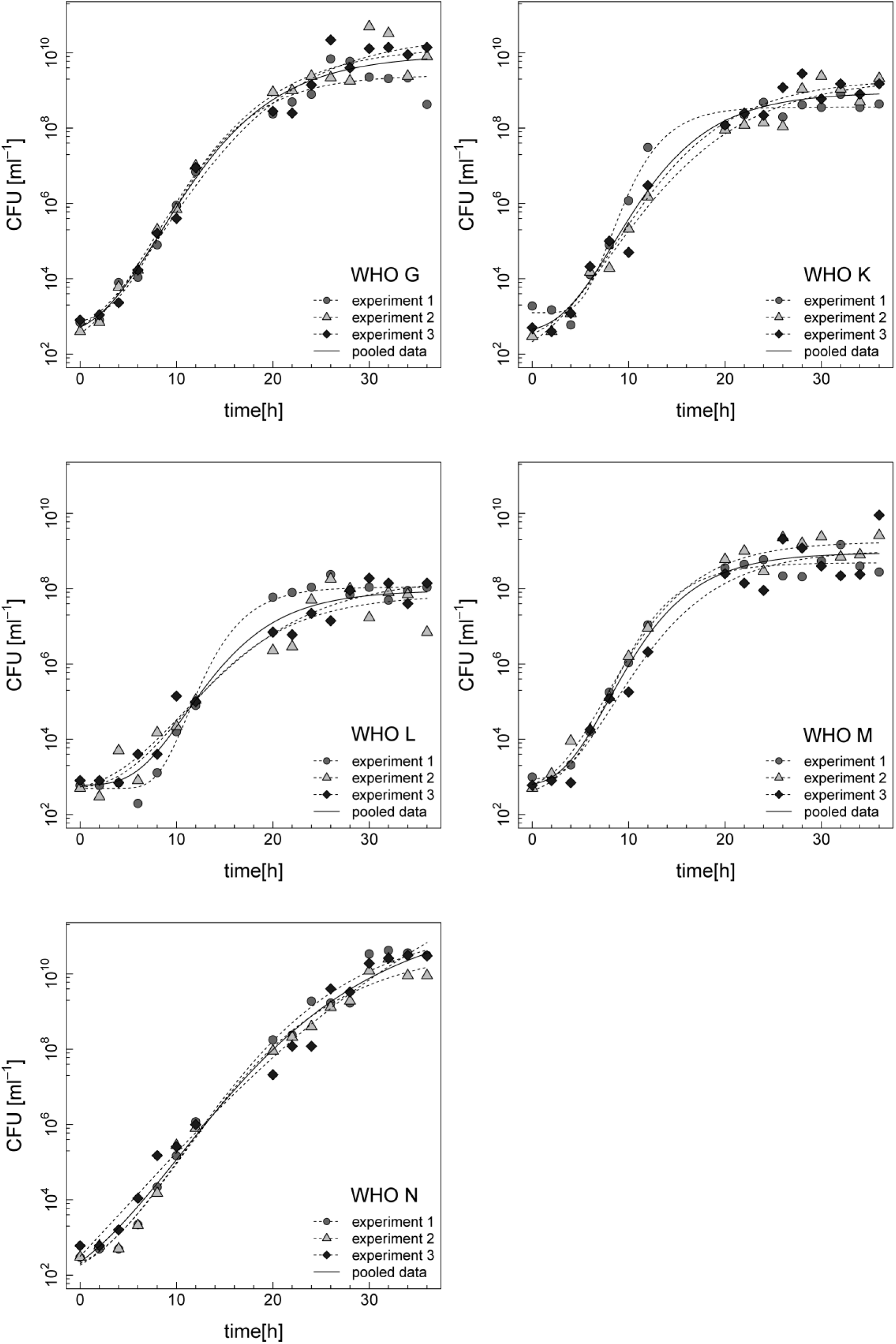
Growth curves for the 2008 WHO *Neisseria gonorrhoeae* reference strains WHO G (A), WHO K (B), WHO L (C), WHO M (D), WHO N (E). Data from three independent experiments are shown. CFU/ml for each time-point are shown in circles (experiment 1), triangles (experiment 2) and diamonds (experiment 3). A Gompertz growth model (1, 2) was fit to the data from three independent experiments (solid lane, pooled data). Individual fits from each of the experiments are shown as well in dashed lines. Growth rates were estimated in log phase between 2–20 hours (WHO G=0.75 [h^−1^], WHO K=0.72 [h^−1^], WHO L=0.57 [h^−1^], WHO M=0.75 [h^−1^], WHO N=0.70 [h^−1^]). The maximal bacterial density was estimated as upper asymptote of the Gompertz model (WHO G=9.74*10^9^ [CFU/ml], WHO K=1.32*10^9^ [CFU/ml], WHO L=6.57*10^7^ [CFU/ml], WHO M=1.32*10^9^ [CFU/ml], WHO N=5.32*10^11^ [CFU/ml]).

**Table S1:**
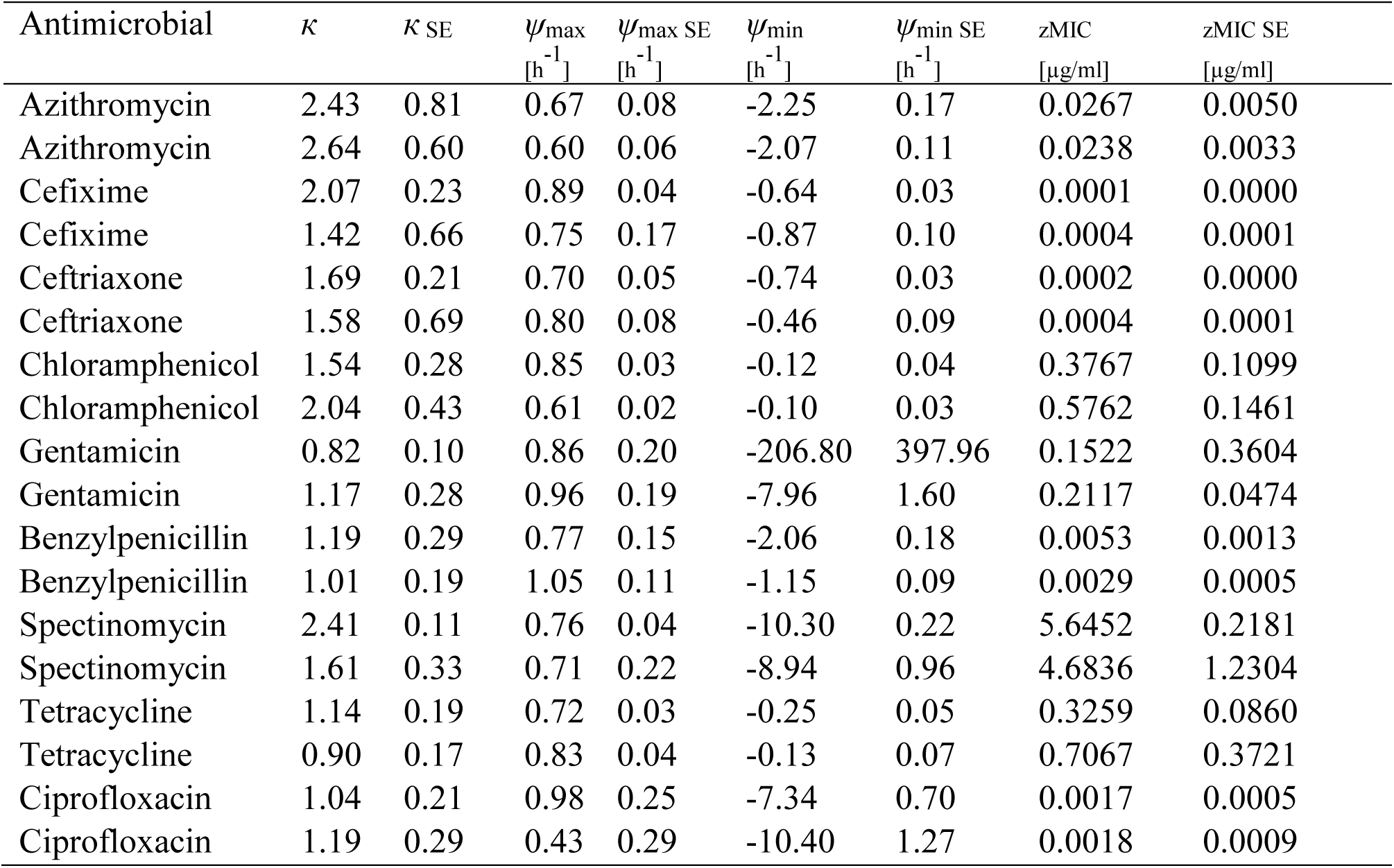
Parameter estimates for nine different antimicrobials in *Neisseria gonorrhoeae* strain DOGK18 and model based standard errors.

**Table S2:** Parameter estimates for ciprofloxacin in five WHO *Neisseria gonorrhoeae* reference strains and DOGK18, and model based standard errors.

| Strain | $K$ | $K$ SE | $\psi_{\max}$ | $\psi_{\max}$ SE | $\psi_{\min}$ | $\psi_{\min}$ SE | zMIC<br>[$\mu\text{g/ml}$] | zMIC SE<br>[$\mu\text{g/ml}$] |
| --- | --- | --- | --- | --- | --- | --- | --- | --- |
| WHO G | 0.69 | 0.14 | 0.86 | 0.12 | -2.22 | 0.35 | 0.0327 | 0.0094 |
| WHO K | 1.47 | 0.29 | 0.75 | 0.03 | -0.74 | 0.18 | 18.3111 | 3.9561 |
| WHO L | 1.56 | 0.27 | 0.89 | 0.06 | -1.02 | 0.10 | 6.3510 | 0.7475 |
| WHO M | 3.49 | 0.77 | 0.62 | 0.05 | -1.18 | 0.04 | 0.2122 | 0.0142 |
| WHO N | 1.58 | 0.24 | 0.81 | 0.04 | -0.60 | 0.04 | 1.7405 | 0.1912 |
| DOGK18 | 1.19 | 0.29 | 0.43 | 0.29 | -10.40 | 1.27 | 0.0018 | 0.0009 |

## REFERENCES

1. Unemo M, Shafer WM. 2014. Antimicrobial resistance in *Neisseria gonorrhoeae* in the 21st Century: past, evolution, and future. Clin Microbiol Rev 27:587–613.

2. Mueller M, de la Peña A, Derendorf H. 2004. Issues in pharmacokinetics and pharmacodynamics of anti-infective agents: kill curves versus MIC. AntimicrobAgents Chemother 48:369–377.

3. Li DRC, Zhu M, Schentag JJ. 2012. Achieving an optimal outcome in the treatment of infections. Clin Pharmacokinet 37:1–16.

4. Regoes RR, Wiuff C, Zappala RM, Garner KN, Baquero F, Levin BR. 2004. Pharmacodynamic functions: a multiparameter approach to the design of antibiotic treatment regimens. Antimicrob Agents Chemother 48:3670–3676.

5. Takei M, Yamaguchi Y, Fukuda H, Yasuda M, Deguchi T. 2005. Cultivation of *Neisseria gonorrhoeae* in liquid media and determination of its in vitro susceptibilities to quinolones. J Clin Microbiol 43:4321–4327.

6. Jeverica S, Golparian D, Hanzelka B, Fowlie AJ, Matičič M, Unemo M. 2014. High in vitro activity of a novel dual bacterial topoisomerase inhibitor of the ATPase activities of GyrB and ParE (VT12-008911) against *Neisseria gonorrhoeae* isolates with various high-level antimicrobial resistance and multidrug resistance. J Antimicrob Chemother 69:1866–1872.

7. Hamilton-Miller JM, Bruzzese T, Nonis A, Shah S. 1996. Comparative anti-gonococcal activity of S-565, a new rifamycin. Int J Antimicrob Agents 7:247–250.

8. Wade JJ, Graver MA. 2007. A fully defined, clear and protein-free liquid medium permitting dense growth of Neisseria gonorrhoeae from very low inocula. FEMS Microbiol Lett 273:35–7.

9. Unemo M, Fasth O, Fredlund H, Limnios A, Tapsall J. 2009. Phenotypic and genetic characterization of the 2008 WHO Neisseria gonorrhoeae reference strain panel intended for global quality assurance and quality control of gonococcal antimicrobial resistance surveillance for public health purposes. J Antimicrob Chemother 63:1142–1151.

10. Chen CY, Nace GW, Irwin PL. 2003. A 6 × 6 drop plate method for simultaneous colony counting and MPN enumeration of Campylobacter jejuni, Listeria monocytogenes, and *Escherichia coli*. J Microbiol Methods 55:475–479.

11. Gagneur J, Neudecker A. 2012. cellGrowth: Fitting cell population growth models. R package version 1.12.0. **Available online**: http://www.bioconductor.org/packages/release/bioc/manuals/cellGrowth/man/cellGrowth.pdf

12. Zwietering MH, Jongenburger I, Rombouts FM, van ’t Riet K. 1990. Modeling of the bacterial growth curve. Appl Environ Microbiol 56:1875–1881.

13. Ritz C, Streibig JC. 2005. Bioassay analysis using R. Stat. Softw. 12, (2005). Available online: http://www.jstatsoft.org/v12/i05/paper

14. R Core Team. 2014. R: A language and environment for statistical computing. R foundation for statistical computing, Vienna, Austria. **Available online**: https://www.r-project.org/

15. LeBel M. 1988. Ciprofloxacin: chemistry, mechanism of action, resistance, antimicrobial spectrum, pharmacokinetics, clinical trials, and adverse reactions. Pharmacotherapy 8:3–33.

16. Borovinskaya MA, Pai RD, Zhang W, Schuwirth BS, Holton JM, Hirokawa G, Kaji H, Kaji A, Cate JHD. 2007. Structural basis for aminoglycoside inhibition of bacterial ribosome recycling. Nat Struct Mol Biol 14:727–732.

17. Borovinskaya MA, Shoji S, Holton JM, Fredrick K, Cate JHD. 2007. A steric block in translation caused by the antibiotic spectinomycin. ACS Chem Biol 2:545–552.

18. Wilson DN. 2009. The A–Z of bacterial translation inhibitors. Crit Rev Biochem Mol Biol 44:393–433.

19. Ward ME. 1977. The bactericidal action of spectinomycin on *Neisseria gonorrhoeae*. J Antimicrob Chemother 3:323–329.

20. Ilina EN, Malakhova MV, Bodoev IN, Oparina NY, Filimonova AV, Govorun VM. 2013. Mutation in ribosomal protein S5 leads to spectinomycin resistance in *Neisseria gonorrhoeae*. Front Microbiol 4:186.

21. Brown LB, Krysiak R, Kamanga G, Mapanje C, Kanyamula H, Banda B, Mhango C, Hoffman M, Kamwendo D, Hobbs M, Hosseinipour MC, Martinson F, Cohen MS, Hoffman IF. 2010. Neisseria gonorrhoeae antimicrobial susceptibility in Lilongwe, Malawi, 2007. Sex Transm Dis 37:169–172.

22. Ross JDC, Lewis DA. 2012. Cephalosporin resistant *Neisseria gonorrhoeae:* time to consider gentamicin? Sex Transm Infect 88:6–8.

23. Dowell D, Kirkcaldy RD. 2012. Effectiveness of gentamicin for gonorrhoea treatment: systematic review and meta-analysis. Sex Transm Infect 88:589–594.

24. Hathorn E, Dhasmana D, Duley L, Ross JD. 2014. The effectiveness of gentamicin in the treatment of *Neisseria gonorrhoeae:* a systematic review. Syst Rev 3:104.

25. Williamson R, Tomasz A. 1985. Inhibition of cell wall synthesis and acylation of the penicillin binding proteins during prolonged exposure of growing *Streptococcus pneumoniae* to benzylpenicillin. Eur J Biochem FEBS 151:475–483.

26. Drusano GL. 2004. Antimicrobial pharmacodynamics: critical interactions of “bug and drug.” Nat Rev Microbiol 2:289–300.

27. Comby S, Flandrois JP, Carret G, Pichat C. 1989. Mathematical modelling of growth of *Escherichia coli* at subinhibitory levels of chloramphenicol or tetracyclines. Res Microbiol 140:243–254.

28. Greulich P, Scott M, Evans MR, Allen RJ. 2015. Growth-dependent bacterial susceptibility to ribosome-targeting antibiotics. Mol Syst Biol 11(1):796.

29. Balaban NQ, Merrin J, Chait R, Kowalik L, Leibler S. 2004. Bacterial persistence as a phenotypic switch. Science 305:1622–1625.

30. Dörr T, Vulić M, Lewis K. 2010. Ciprofloxacin causes persister formation by inducing the TisB toxin in *Escherichia coli*. PLoS Biol 8:e1000317.

31. Feng J, Kessler DA, Ben-Jacob E, Levine H. 2014. Growth feedback as a basis for persister bistability. Proc Natl Acad Sci USA 111:544–549.

32. Maisonneuve E, Gerdes K. 2014. Molecular mechanisms underlying bacterial persisters. Cell 157:539–548.

33. Lewis K. 2007. Persister cells, dormancy and infectious disease. Nat Rev Microbiol 5:48–56.

34. Kint CI, Verstraeten N, Fauvart M, Michiels J. 2012. New-found fundamentals of bacterial persistence. Trends Microbiol 20:577–585.

35. Fauvart M, De Groote VN, Michiels J. 2011. Role of persister cells in chronic infections: clinical relevance and perspectives on anti-persister therapies. J Med Microbiol 60:699–709.

36. Conlon BP. 2014. Staphylococcus aureus chronic and relapsing infections: Evidence of a role for persister cells: An investigation of persister cells, their formation and their role in *S. aureus* disease. BioEssays News Rev Mol Cell Dev Biol 36:991-.

37. Jaffe HW, Schroeter AL, Reynolds GH, Zaidi AA, Martin JE, Thayer JD. 1979. Pharmacokinetic determinants of penicillin cure of gonococcal urethritis. Antimicrob Agents Chemother 15:587–591.

38. Deguchi T, Yasuda M, Yokoi S, Ishida K-I, Ito M, Ishihara S, Minamidate K, Harada Y, Tei K, Kojima K, Tamaki M, Maeda S-I. 2003. Treatment of uncomplicated gonococcal urethritis by double-dosing of 200 mg cefixime at a 6-h interval. J Infect Chemother Off J Jpn Soc Chemother 9:35–39.

39. Chisholm SA, Mouton JW, Lewis DA, Nichols T, Ison CA, Livermore DM. 2010. Cephalosporin MIC creep among gonococci: time for a pharmacodynamic rethink? J Antimicrob Chemother 65:2141–8.

40. Faulkner RD, Bohaychuk W, Lanc RA, Haynes JD, Desjardins RE, Yacobi A, Silber BM. 1988. Pharmacokinetics of cefixime in the young and elderly. J Antimicrob Chemother 21:787–794.

41. Meyers BR, Srulevitch ES, Jacobson J, Hirschman SZ. 1983. Crossover study of the pharmacokinetics of ceftriaxone administered intravenously or intramuscularly to healthy volunteers. Antimicrob Agents Chemother 24:812–814.

42. Drusano GL. 2007. Pharmacokinetics and pharmacodynamics of antimicrobials. Clin Infect Dis Off Publ Infect Dis Soc Am 45 Suppl 1:S89–95.

43. Levison ME, Levison JH. 2009. Pharmacokinetics and pharmacodynamics of antibacterial agents. Infect Dis Clin North Am 23:791–815, vii.

44. Craig WA. 1998. Pharmacokinetic/pharmacodynamic parameters: rationale for antibacterial dosing of mice and men. Clin Infect Dis Off Publ Infect Dis Soc Am 26:1–10; quiz 11-12.

45. Frimodt-Møller N. 2002. How predictive is PK/PD for antibacterial agents? Int J Antimicrob Agents 19:333–339.

46. Dorfman MS, Wagner RS, Jamison T, Bell B, Stroman DW. 2008. The pharmacodynamic properties of azithromycin in a kinetics-of-kill model and implications for bacterial conjunctivitis treatment. Adv Ther 25:208–217.

47. Imamura Y, Higashiyama Y, Tomono K, Izumikawa K, Yanagihara K, Ohno H, Miyazaki Y, Hirakata Y, Mizuta Y, Kadota J, Iglewski BH, Kohno S. 2005. Azithromycin Exhibits Bactericidal Effects on Pseudomonas aeruginosa through Interaction with the Outer Membrane. Antimicrob Agents Chemother 49:1377–1380.

## REFERENCES

1. Gagneur J, Neudecker A. 2012. cellGrowth: Fitting cell population growth models. R package version 1.12.0. **Available online**: http://www.bioconductor.org/packages/release/bioc/manuals/cellGrowth/man/cellGrowth.pdf

2. Zwietering MH, Jongenburger I, Rombouts FM, van ’t Riet K. 1990. Modeling of the bacterial growth curve. Appl Environ Microbiol 56:1875–1881.

